# “Human platelet lysate derived extracellular vesicles enhance angiogenesis through miR-126”

**DOI:** 10.1101/2022.05.10.491341

**Authors:** Antonella Bordin, Maila Chirivì, Francesca Pagano, Marika Milan, Marco Iuliano, Eleonora Scaccia, Orazio Fortunato, Giorgio Mangino, Xhulio Dhori, Elisabetta De Marinis, Alessandra D’Amico, Selenia Miglietta, Vittorio Picchio, Roberto Rizzi, Giovanna Romeo, Fabio Pulcinelli, Isotta Chimenti, Giacomo Frati, Elena De Falco

## Abstract

**Objectives:** extracellular vesicles (EVs) are key biological mediators of several physiological functions within the cell microenvironment. Platelets are the most abundant source of EVs in the blood. Similarly, platelet lysate (PL), the best platelet derivative and angiogenic performer for regenerative purposes, is enriched of EVs, but their role is still too poorly discovered to be suitably exploited. Here we explored the contribution of the EVs in PL, by investigating the angiogenic features extrapolated from that possessed by PL.

**Methods:** we tested angiogenic ability and molecular cargo in 3D bioprinted models and by RNA sequencing analysis of PL-derived EVs.

**Results:** a subset of small vesicles is highly represented in PL. The EVs do not retain aggregation ability, preserving a low redox state in HUVEC and increasing the angiogenic tubularly-like structures in 3D endothelial bioprinted constructs. EVs resembled the miRNome profile of PL, mainly enriched of small RNAs and a high amount of miR-126, the most abundant angiogenic miRNA in platelets. The transfer of miR-126 by EVs in HUVEC after the *in vitro* inhibition of the endogenous form, restored angiogenesis, without involving VEGF as downstream target in this system.

**Conclusions:** PL is a biological source of available EVs with angiogenic effects involving a miRNAs-based cargo. These properties can be exploited for targeted molecular/biological manipulation of PL, by potentially developing a product exclusively manufactured of EVs.

## INTRODUCTION

Beyond haemostasis and thrombosis, platelets have been also described as main regulators of angiogenesis, a key process for tissue regeneration and repair outcome of vascular insults or wound healing and based on the activation of endothelial proliferation, sprouting and organization into functional tubules^1–3^. The emerging role of platelets to act as inflammatory/immune effectors and to enhance angiogenesis, stems from their intrinsic physiological role to interact with the endothelium during vascular damage, preserving integrity and vessel homeostasis^4^.

Platelets exhibit a unique secretory profile of multiple combined factors with a dual pro- and anti-angiogenic role. Among them, we could list growth factors, cytokines, microRNAs, small soluble molecules and proteins, including those related to cytoskeleton, adhesion, inflammation, and extracellular matrix interaction^5–8^. This balanced combination of mediators, is mainly contained in plasma membranes delimited nanoparticles, named extracellular vesicles (EVs). These latter, now conceived as signalosomes and biological vectors of heterogeneous size and composition, are released upon platelets activation, interacting with the microenvironment^9,10^.

Platelets represent the most abundant source of EVs of different dimension and quantity in the systemic circulation^11–14^depending on multiple variables including age, physiological states, and lifestyle habits^15,16^. EVs mirror the haemostatic properties of platelets^17^, by exerting both anti- and pro-coagulant effects according to the subpopulation of EVs involved^11,18,19^. Hence, their circulating levels can act as predictive biomarkers of haemostatic and inflammatory disorders^20^. Patients with metabolic syndrome, myocardial infarction, atherosclerosis, ischemia, or inflammatory diseases exhibit higher levels of circulating EVs, because of activated platelets^21–28^, suggesting their relevance to mediate pathogenetic effects beyond their physiological role. On the other hand, platelets-derived EVs have been also demonstrated to regulate angiogenesis when released at the site of endothelial sprouts ^29^ and secretion of VEGF^30^, or to transfer proliferative and survival biological information to the endothelium^30–33^. Platelet-derived EVs can modulate the vascular tone as shown in rabbit models^30^, or even attenuate blood pressure in preeclampsia women, by stimulating the inducible nitric oxide synthase in endothelial cells^34^, consistently with a beneficial functional role on the vasculature.

Similarly to their parental cells, EVs would also act indirectly in a paracrine fashion on the intercellular network as cargos of cytokines and decoy proteins locally released in the microenvironment. For instance, endothelial progenitor cell-mediated angiogenesis^1,3^and tissue engraftment is enhanced by EVs of platelets origin through the activation of specific targets (MMP-2, 9, PI3-Kinase, ERK) or after transfer of specific soluble platelet receptors and activation of integrins on the endothelial surface^30,35–37^. The preconditioning of bone marrow mesenchymal stem cells with platelet-derived EVs has demonstrated their effective capacity of boosting the biological potency and vascular effects of the stromal fraction^38^. Strong evidence of their ability to support angiogenesis has been also observed in myocardial infarction and cerebral ischemia after *in vivo* direct injection, when platelets, activated by thrombin, release EVs^39,40^. Moreover, the key contribution of platelet-derived EVs in supporting the angiogenic profile of cancer invasion and metastasis parallel to clinical thrombotic complications has been strongly highlighted^41–43^.

Based on these studies, evidence that platelets and platelet-derived biological products can trigger angiogenic programs in endothelial cells has encouraged a better understanding of their potential therapeutic use for those regenerative-based applications where the restoration or the enhancement of angiogenesis represents the clinical goal. Accordingly, parallel to the investigations regarding the key involvement of platelets in regulating angiogenesis, it has been clearly demonstrated that platelet-derived clinical preparations (*i.e*. platelet-rich plasma, and gels) are similarly able to boost and reflect the angiogenic properties of platelets. Particular attention has been dedicated to platelet lysate (PL), considered as the gold preparation concentrate derived from platelets, and whose clinical efficacy is currently considered superior to other platelet-derived formulations^44^. The employment of PL, alone or even in combination with different sources of stem cells, has shown to enhance blood perfusion in peripheral artery diseases^45^, to heal difficult wounds, to sustain stromal proliferation, epithelization, angiogenesis, and to prime cardiovascular differentiation^1–3,46–49^. The angiogenic capacity of PL is ascribable to the plethora of highly concentrated factors in this hemoderivative. When PL is manufactured, platelets are repeatedly lysed, therefore enriching the preparations with vesicles and granules, representing a primary source of angiogenic EVs. So far, the vast majority of studies has only explored the effects of vesicles of different origin (*i.e*. from MSCs, fibroblasts, lymphocytes) after treatment with PL, or EVs released by intact and activated platelets^43^. Only a couple of studies have described the presence of exosomes in platelet-derived clinical formulations but as effectors of the osteogenic differentiation on MSCs or with neuroregenerative capacities^38,50^. Thus, the wide range of mechanisms by which PL-derived EVs might regulate angiogenesis still needs to be fully addressed.

This study investigates form a biological and molecular standpoint the role of EVs in relation to PL formulations regarding the ability to mediate angiogenesis by endothelial cells.

## METHODS

### Isolation of Extracellular vesicles from Platelet lysate-based preparations

Extracellular vesicles were isolated from PL-based preparations^2,46–49,51^ by sequentially ultracentrifugation (500 rcf for 10 minutes; 2000 rcf for 10 minutes; 100,000 rcf for 1 hour). EV pellet was then resuspended to reconstitute the initial PL volume and subsequently used at 5, 10% or 20% in medium for the experiments. For FACS analysis and uptake assay in HUVEC, 100μl of EVs were stained for 10 minutes at 37°C with 5 μM 5(6)-CFDA-SE [5-(and-6)-Carboxyfluorescein Diacetate, Succinimidyl Ester (CFSE, Invitrogen/Thermo Fischer Scientific) according to the manufacturer’s instructions. Excess dye was removed using Exosome Spin Columns (Thermo Fischer Scientific) following manufacturer’s recommendations.

### Nanotracking Analysis of extracellular vesicles in PL

Nanoparticles tracking analysis in terms of size distribution and concentration was performed on PL using a NanoSight NS300 instrument (Malvern Panalytical). Five 30 seconds videos were recorded for each sample with a camera level set at 15/16 and a detection threshold set between 5 and 7. The EVs concentration and size distribution were subsequently analysed with NTA 3.2 software.

### Western Blot

EV were resuspended in a RIPA buffer and phosphatase inhibitor cocktail. Proteins (10μl solubilized in 2X Laemmli/20% of 2-mercaptoethanol) were separated by SDS-PAGE on 10% polyacrylamide gel (Bio-Rad, Hercules). Subsequently, the membrane was blocked with 5% (w/v) milk and the membranes incubated at 4°C overnight with rabbit Anti-Annexin A1/ANXA1 antibody monoclonal antibody [EPR19342]-BSA and Azide free (1: 25000, Abcam, Cat. N. ab222398), ALIX (1:1000, Biorbyt, Cat. N. orb235075), CD9 (1:500, Abcam, ab186429), calnexin (1:1000, Santa Cruz, Cat. N. sc-46669, TSG101 (4A10) (1:500, Invitrogen Cat. N. #MA1-23296) After incubation, the membranes were incubated with secondary anti-rabbit antibody (Cell Signaling, 1:10000) and the immune complexes thus formed were detected by enhanced substrate chemiluminescence. Densitometric detection of the bands was performed by Chemidoc (Biorad).

### Cell culture, transfection and treatment with AntagomiR-126

Human umbilical vein endothelial cells (HUVEC) were cultured between passage 3 and 6 in EGM-2 complete medium (Lonza)^3,52–55^. Fluorescein-conjugated Antagomirs were used for quantification of Antagomir or control incorporation and detected by flow cytometry. For AntagomiR-126 transfection, cells were plated at a density of 3.5×10^4^/24 wells with EGM-2 without gentamicin. A mix composed of 25 picomoles LNA_126 (miRCURY LNA miRNA Inhibitor (5) - 3 ‘Fam, Cat. N. 339121, Qiagen) or 25 picomoles Control (miRCURY LNA miRNA Inhibitor Control (5) -No Modification Fam, Cat N. 339126. Qiagen) in Optimem reduced serum media and lipofectamine (1μl/100 μl Optimem, RNAiMAX, Invitrogen, Cat. N. 56531) was added to the HUVEC and incubated overnight. The next day, the medium was removed, and new EGM-2 was added to the cells for up to 24 and 48 hours of total transfection. To verify transfection, HUVEC were analyzed by flow cytometry detecting FAM fluorescence.

Cytometric Analysis, Transmission electron microscopy, Matrigel assay, and Immunofluorescence on HUVEC in 3D-bioprinting constructs, Platelets aggregation, determination of soluble human prothrombin Fragment 1+2 and quantification of Peroxide hydrogen and Molecular Biology

See supplementary methods

### Statistics

Statistical Analysis was performed by GraphPad PRISM 5 software. Student’s t-test and One-way ANOVA (Bonferroni correction) were used to compare the difference between the control and groups. A p value <0.05 was considered as significant. Data were presented as mean ±standard error unless specified. Additional information on statistics and confidence intervals have been reported in the corresponding sections above and in figure legends.

## RESULTS

We investigated in detail the EV content and characteristics of human PL preparations. In order to assess the concentration and absolute size distribution^56^, we measured the EVs by Nanoparticle Tracking Analysis (NTA) in eight different batches of PL. Results showed that PL-based preparations contain a very high concentration of EVs with a mean of 1.84×10^13^particles/ml, with the EV size mode of 123.37±7.02 nm (Fig. 1a-b). Among all distinctive subclasses of EVs, the most representative group in terms of concentration was the 50-200 nm subset, corresponding to small microvesicles including exosomes^57^ (Figure1c-d, p<0.001 *vs* all subsets). Occasionally, EVs of <50nm size were found, but not in all batches. Following international guidelines^58,59^, EVs were further characterized by evaluating their morphology and phenotype by transmission electronic microscopy (TEM), and western blot for the recommended universal markers. TEM analysis confirmed that PL preparations contained EVs with heterogeneous but small dimension, roundish morphology, and electron-dense features, suggesting a significant cargo function (Fig. 1e). According to the Minimal Information for Studies of Extracellular Vesicles (“MISEV”) guidelines^58^, TEM was qualitatively implemented by both Western Blot showing the positive expression of the proteins ALIX, CD9, Annexin A1, TSG101 (cytosolic, membrane and marker of biogenesis of EV) in PL-derived EVs^60,61^ and negative expression for calnexin, therefore suggesting the absence of nonEV structures in the preparation of EVs^62^ (Fig. 1f). The characterization of EVs was further verified by cytofluorimetry. The FACS Analysis confirmed the expression of CD41, a main marker of platelet origin of EVs (49.92%±5.22, also known as glycoprotein IIb possessing a critical role in modulating platelet aggregation^63^), but also the negative expression for calnexin, (Fig. 1g).

**FIGURE 1.**
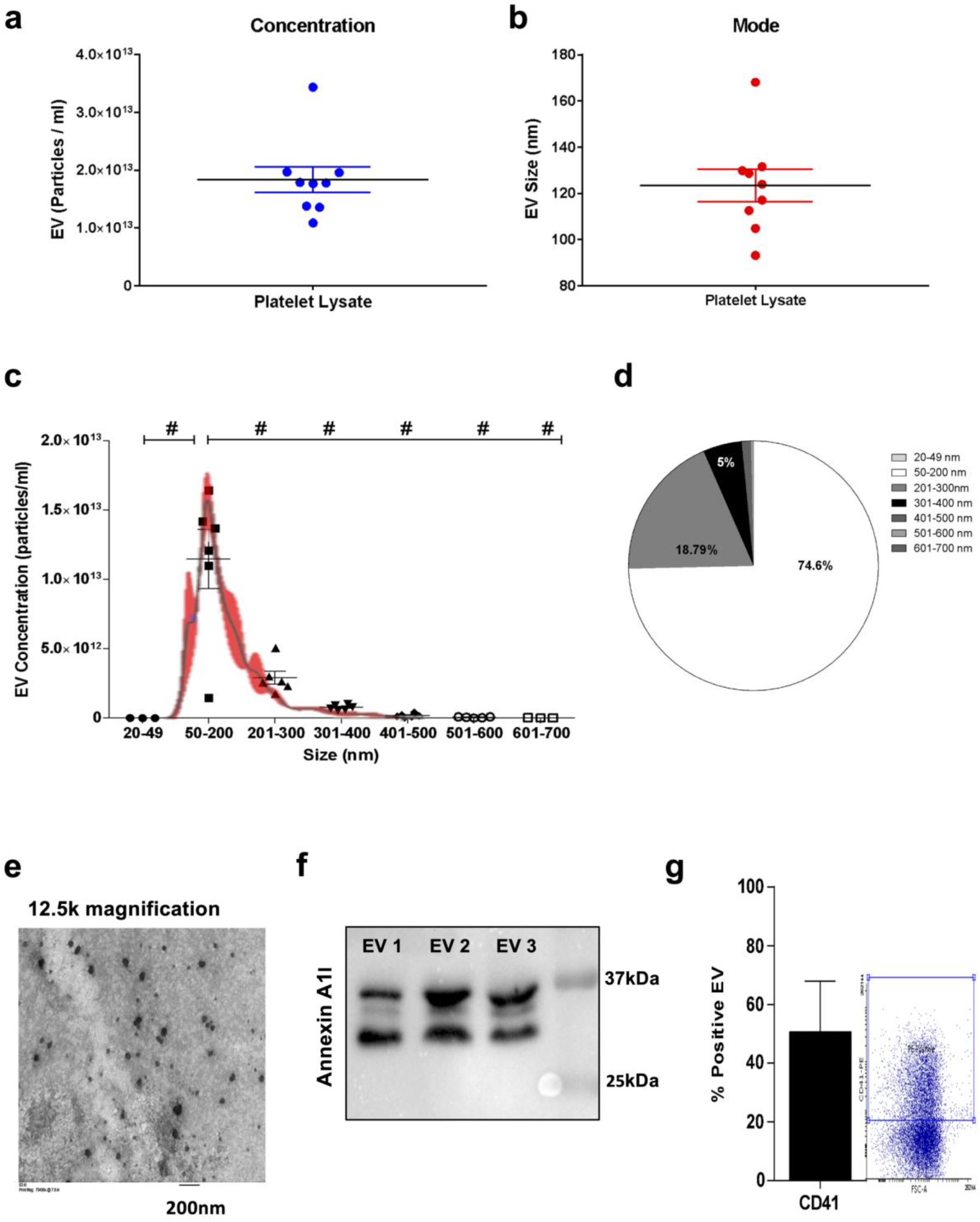
Characterization of Extracellular Vesicles in Platelet Lysate. (A) Concentration measurement and (B) size Mode from Nanoparticle Tracking Analysis (NTA) of EVs derived by 9 batches of PL. (C) NTA graph and (D) pie chart display the EVs size distributions. # p<0.001. (E) Representative transmission electron microscopy images of EVs within PL-based preparations. Scale bar (200nm) and magnification (20000X) are displayed. (F) Western Blot analysis of ALIX, calnexin, TSG101, Annexin A1 and CD9 of 3 EV batches isolated by ultracentrifugation from PL. (G) Representative histogram of the Flow Cytometry displaying the relative percentage of EVs in the graph below for CD41, platelet marker. N=3 lots of EVs were tested.

In order to discriminate the biological effects of EVs from the whole PL, we isolated the EVs according to methodological standardized guidelines by high-speed ultracentrifugation^64,65^. Afterwards, we investigated whether EVs may convey haemostatic properties, such as aggregation and procoagulant abilities, that are two key physiological properties exerted by platelets, but also reported for EVs^66^. We stimulated platelets of healthy subjects with increasing percentages of EVs (5, 10 and 20%). Platelet lysate (10 and 20%) and Collagen were used as biological and positive reference, respectively. Additionally, the quantification of the soluble fragment 1+2 of prothrombin (F1+2) was employed to test the coagulation property of EVs. Results showed that neither increasing concentration of EVs nor PL were able to induce aggregation compared to collagen (Fig. 2a-b). A similar amount of F1+2 among samples was detected (comparable to physiological soluble levels in the human plasma), with no statistically significant differences (Fig. 2c).

**FIGURE 2.**
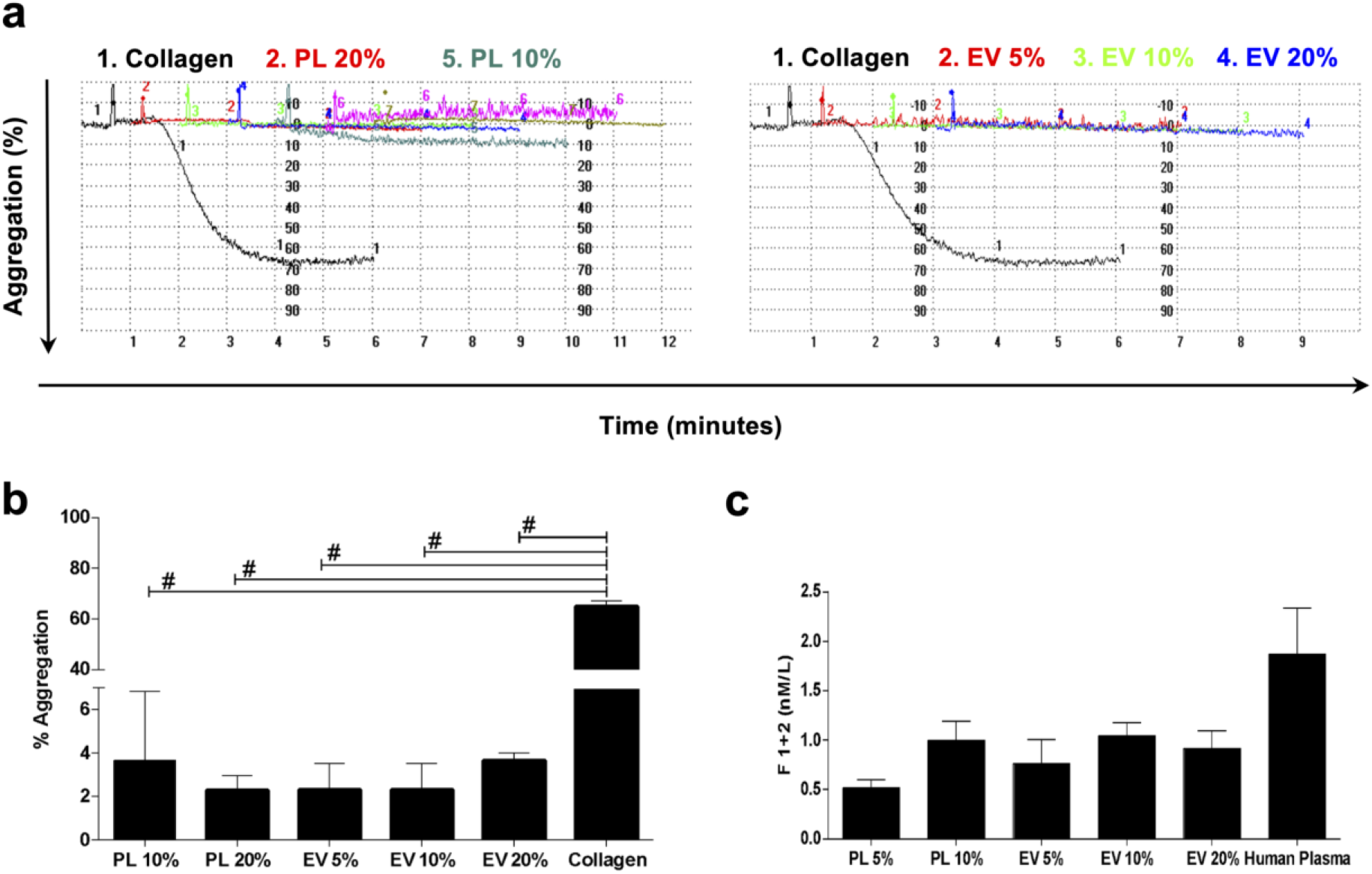
Hemostatic properties of EVs derived from PL. (A) Representative plots of the aggregometer displaying both PL and EVs at different percentage and (B) relative analysis showing no aggregation compared to collagen used as positive reference. # p<0.001 (C) Immunoassay of the F1+2 assaying the coagulative ability of both PL and EVs compared with Human Plasma with no significant difference.

As one of the most significant bioactive properties of PL is the ability to induce angiogenesis^2,46–49^, we investigated the contribution of EVs to the angiogenesis stimuli mediated by PL. We isolated and labelled the EVs with the green fluorescent dye CFSE. Afterwards, human umbilical vein endothelial cells (HUVECs) were stimulated for 24 hours with the EV preparation (10% v/v, corresponding to the same PL volume in percentage routinely employed in cell culture^47–49,51^). HUVECs were able to uptake EVs, as demonstrated by the presence of green fluorescent dots visible in the cytoplasm (Fig. 3a). When HUVECs were subjected to the *in vitro* angiogenesis Matrigel assay at increasing concentrations of EVs (5, 10, 20%), we found that 10% EVs was the optimal percentage to significantly enhance the number of closed loops (Fig. 3b-c) compared to the negative control (EBM, non-supplemented Endothelial basal media, p=0.0038). Platelets lysate and EGM-2 were used as positive angiogenic inducers. To corroborate this observation, we employed a 3D bioprinting-based approach by encapsulating HUVECs in a gelatin/methacrylamide (GelMA) bioink, to evaluate their ability to induce the generation of vessel-like structures in a more physiologically suitable 3D microenvironment in presence of 10% EVs (the best performer in the matrigel assay). The confocal microscopy analysis showed that endothelial cells were able to colonize the bioprinted construct after treatment with both PL and PL-derived EVs. Coherent with the observed spatial distribution, the proportion of the endothelial area (defined as CD31^+^/vWF^+^), corresponding to the organization of HUVECs in 3D tubular structures, was significantly higher with PL-derived EV and PL treatments, compared to EBM control (Fig. 3d-e, p<0.05 *vs* 10% EVs and 10% PL).

**FIGURE 3.**
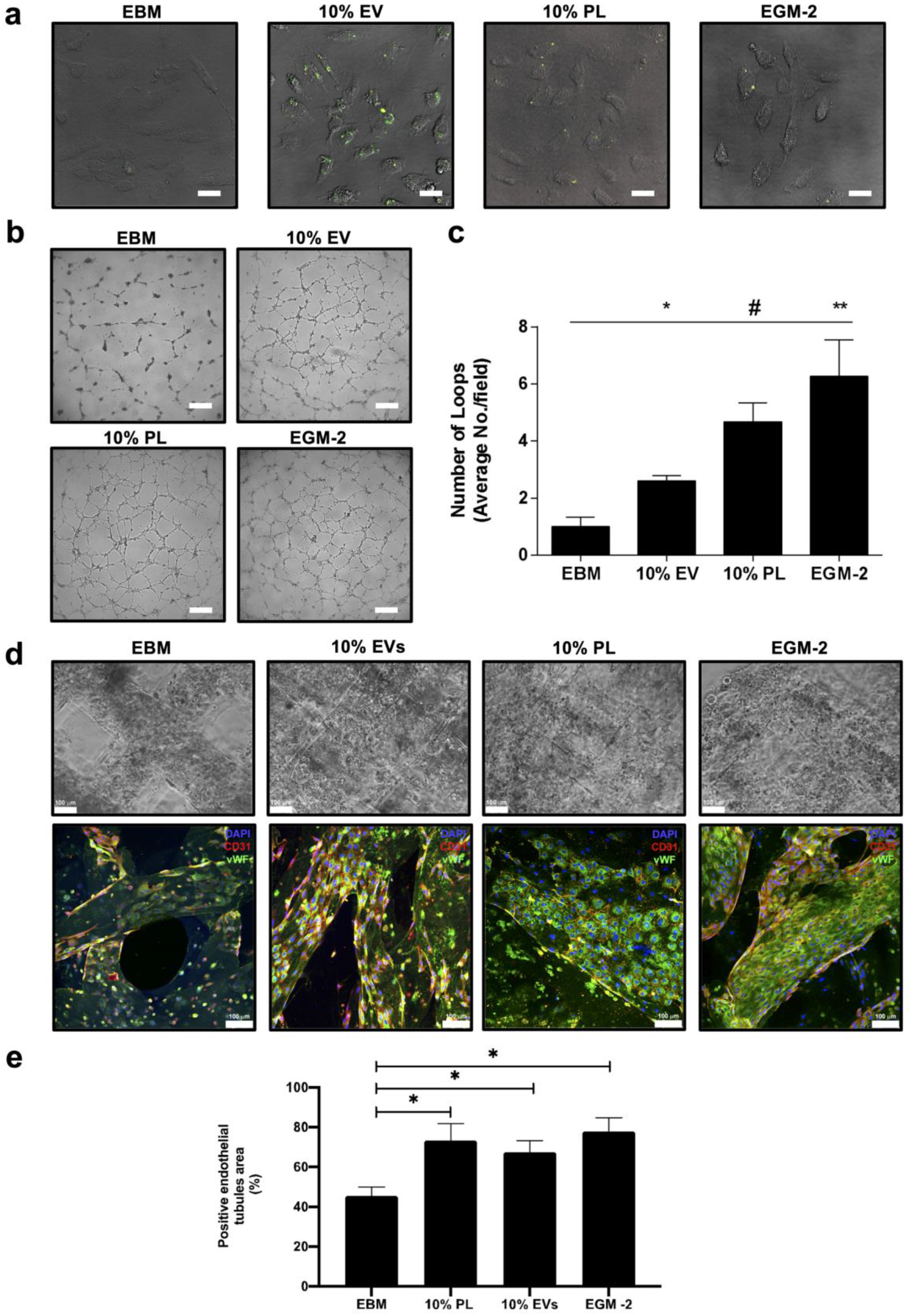
Angiogenic effects of PL-derived EVs in endothelial cells. (A) Merged images of optical and fluorescent microscopy showing HUVEC uptaking after 24 hours the CSFE-labeled isolated EVs in different experimental conditions (EBM and EGM-2 negative and positive controls, respectively). White scale bar, 50μm. (B) Representative images of capillary-like structures from Matrigel assay and (C) quantitative analysis of number of formed loops with different percentage of EVs (5, 10 and 20%). Magnification 4X. White scale bar, 200μm. *p<0.05, **p<0.01, # p<0.001. One way ANOVA test was applied. A range of N=3-12 experiments was performed (D) Confocal microscopy images of 3D *in vitro* bioprinted HUVEC (Bright and fluorescent images), displaying the formation of 3D-tubules angiogenic structures. DAPI, CD31 and vWF stain blue, red and green, respectively. White scale bar, 100μm. (E) Analysis of the immunofluorescence indicating the percentage of the positive 3D endothelial tubules area in the different conditions. *p<0.05.

Several studies have demonstrated the modulation of the redox status in cells exposed to intact platelet-derived EVs^67^. Thus, we investigated the levels of hydrogen peroxide in the conditioned media of HUVECs collected after 24 hours of treatment with EVs. Results showed a lower release of hydrogen peroxide after treatment with 10% EVs compared to PL (p=0 .03, Fig. 4a). We observed that the treatment with all percentages of EVs were able to maintain very low amounts of H2O2 in the media as both controls (EBM and EGM-2). However, the 10% EVs reveals as the optimal anti-oxidant stimulation respect to 10% PL (p<0.05). This result was also coherent with the lowest expression level of the NADPH isoform Nox4 (the main and specific isoform responsible for the direct production of hydrogen peroxide by endothelial cells^52,53,68–70^) after stimulation with 10% EVs among the 3 concentrations of EVs (p<0.05, Fig-4b). The Nox4 mRNA levels in presence of 10% EVs were similarly downregulated as PL and EGM-2 respect to EBM (p<0.05).

**FIGURE 4.**
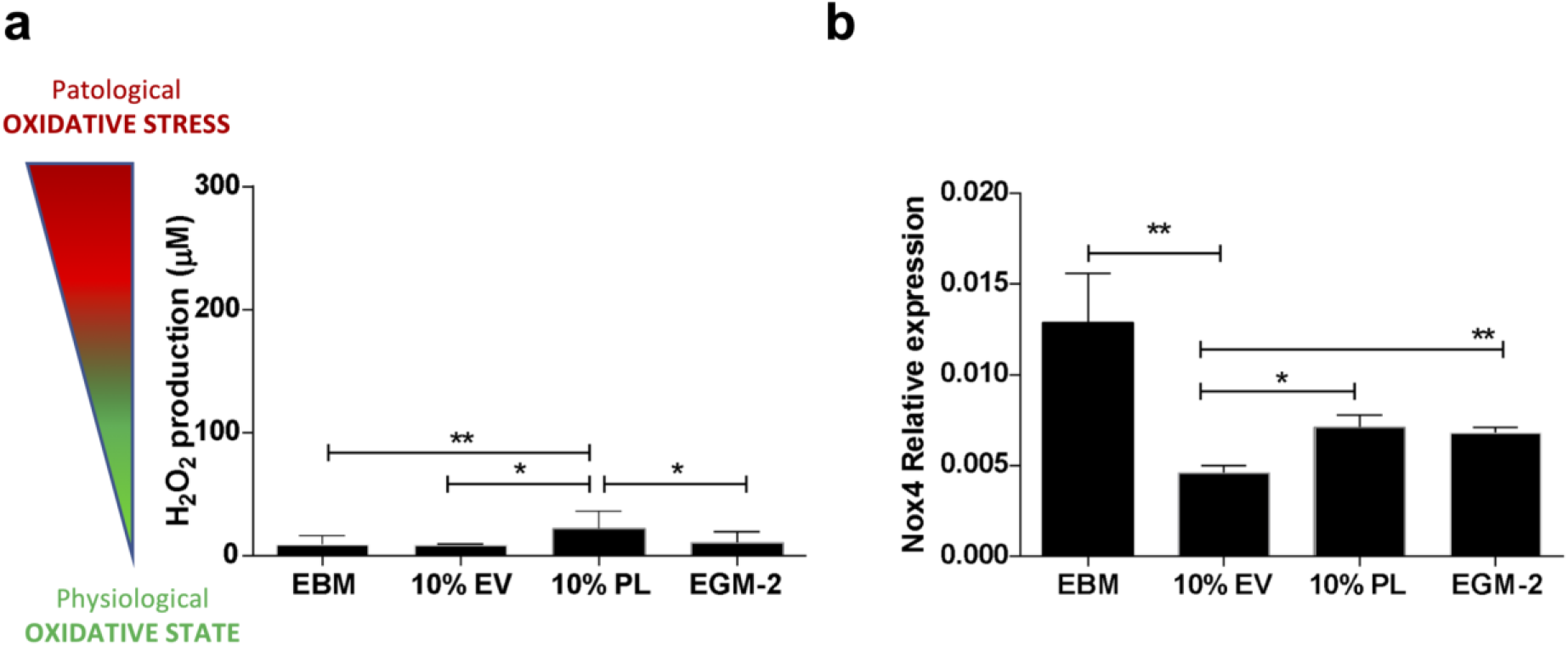
Analysis of the oxidative states of endothelial cells after treatment with different percentage of PL-derived EVs (5, 10, 20%). (A) Hydrogen Peroxide (H2O2) measurement of HUVEC-derived condition media in all experimental conditions. The cartoon on the left of the grap^h^ shows the wide concentration range of H_2_O_2_ from physiological states (≤100μM) to oxidative stress (≥300μM). *p<0.05, **p<0.01. One way ANOVA test was applied. A range of N=3-16 experiments was performed. (B) Relative expression of the NADPH oxidase isoform 4 (NOX4) assayed by Real Time PCR and downregulated in HUVEC after treatment with 10% EVs respect to EBM and all other percentage of EVs (5 and 20%). The effect is also and comparable to that exerted by 10% PL and EGM-2. *p<0.05. One way ANOVA test was applied. A range of N=3-11 experiments was performed.

Some key functions of platelets, such as aggregation, activation, and angiogenesis, are known to be mediated by miRNAs released by platelets in response to a wide range of stimuli, both physiological and pathological^71,72^. This ability can also be mediated by EVs, since they are known to transfer information to target cells through miRNAs^73^, and therefore to determine diverse biological effects in relation to the cargo within the vesicles. With these premises, we hypothesized the presence of miRNAs in PL based formulations and assessed this by analysing the miRNA profile of two different batches of PL for a total of 4 replicates. Results showed that the majority of the small RNA content in PL is represented by miRNAs (43%), followed by Y RNAs (17%), antisense RNAs (10%), and lincRNAs (8%) (Fig. 5a). A miscellaneous group is also represented (22%). After applying a cut-off of >10 copies in all analyzed batches of PL (average count among the 4 PL samples; Table 1), the identified miRNAs clustered into 3 macro-groups based on their expression levels (low, medium and high) when analyzed by heatmap with hierarchical clustering (Fig. 5b).

**FIGURE 5.**
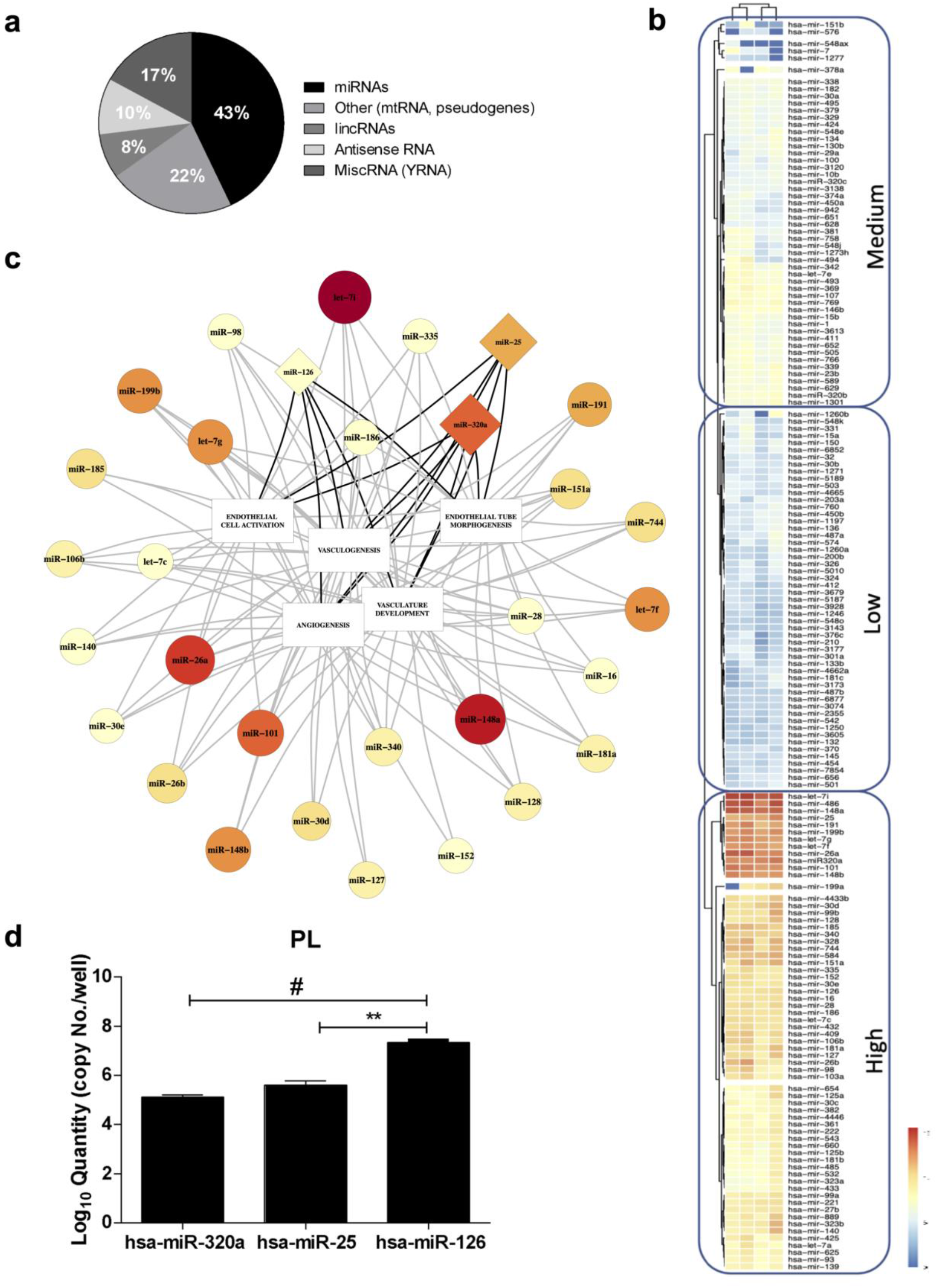
MiRNome characterization of Platelet Lysate by Small RNA sequencing. (A) Small RNA percentage distribution in PL (a total of 4 replicates). (B) The heatmap has been obtained by including highly and consistently expressed miRNAs among the 4 analyzed replicates with a cut-off of >10 copies. The red-yellow and the blue range color indicate miRNAs with high and low average copy count, respectively. (C) Integration of the biological function with the expression of miRNAs in PL using a linkage group intertwined with 5 GO terms. Both the dimension of the box (circle/triangle) and the color range reflect the expression levels as in the heatmap. (D) Quantitative PCR of hsa-miR-hsa-320a, hsa-miR-25 and hsa-miR-126 have been employed to quantitatively validate the miRNome of PL. These 3 miRNAs are represented in the triangle box in C.

As a further selective step, we set a threshold for miRNAs with over 1000 reads, thus obtaining a shortlist of 39 miRNAs (Table 1), which underwent a bioinformatic top-down analysis on the mirPath v.3 online tool (reverse search). The Gene Ontology (GO) terms selection was performed according to the function of interest observed for PL treatment on endothelial cells, which is angiogenesis. We interrogated the database by employing five GO, including the first two with the highest hierarchy for processes of capillary and vascular formation, in particular: VASCULOGENESIS (GO_0001570), ANGIOGENESIS (GO_0001525), ENDOTHELIAL CELL ACTIVATION (GO_0042118), VASCULAR DEVELOPMENT (GO_0001944) and ENDOTHELIAL TUBE MORPHOGENESIS (GO_0061154).

After intersecting each GO term with the top 39 miRNA shortlist, we extrapolated potential eligible candidates for the abovementioned roles. We found that the highest overlap was with the angiogenesis gene list as displayed in the function-expression interaction network that we generated by software (Fig. 5c). Afterwards, we compared the results and shortlisted a main group of 31 miRNAs and a further subset of 11 miRNAs correlating with two and all five GO categories, respectively, where 3 miRNAs (hsa-miR-320a, hsa-miR-25 and hsa -miR-126) were selected to quantitatively validate the seq data by Realtime PCR (Table 2). Notably, miR-126 is a key regulator of angiogenesis and is known as the angio-miRNA and one of the most abundant and specific miR to endothelial cells, human platelets, and platelet-derived vesicles^74^. Notably, the qPCR data accurately highlighted that miR-126 is the most abundant miRNA in PL, compared to miR-320a and miR-25 (Fig. 5d, p<0.01 and p>0.001).

Next, we validated the role of miR-126 in the biological effects of PL and EVs on HUVECs, by comparing the sole stimuli that was the media supplemented with 10% PL or 5, 10 and 20% EVs (with volumes and dilutions adjusted to correspond to 10% PL). Although an increasing but not statistically significant trend of miR-126 was observed among the percentages of EVs, Realtime PCR testing confirmed the equivalent content of miR-126 in all media recipes (Fig. 6a). The treatment with 10% EVs was the only able to significantly upregulate the levels of intracellular miR-126 in endothelial cells compared to the negative control (Fig. 6b, p<0.05). We sought to verify whether EV-miR-126 could play a direct role in mediating angiogenesis ascribable to the transferring of this miRNA through EVs. To this aim, we transfected HUVECs with the antagomir-126 (LNA-126) to rule out endogenous contribution^75^. Then, we stimulated the cells with 10% EVs or PL. Results showed that HUVECs were efficiently transfected by both the LNA-126 and Control, as shown by flow cytometry analysis (supplementary Fig. 1a, 84.84±0.4% positive cells with LNA-126, and 78.5±5.15% with Control). Moreover, the copy number of miR-126 in HUVEC, quantified by droplet digital PCR, was significantly decreased at the lowest levels 24 hours after treatment with the LNA-126, compared to both untreated and transfection controls (supplementary Fig. 1b, p<0.01 and p<0.05, respectively), while at 48h the levels had increased again.

**FIGURE 6.**
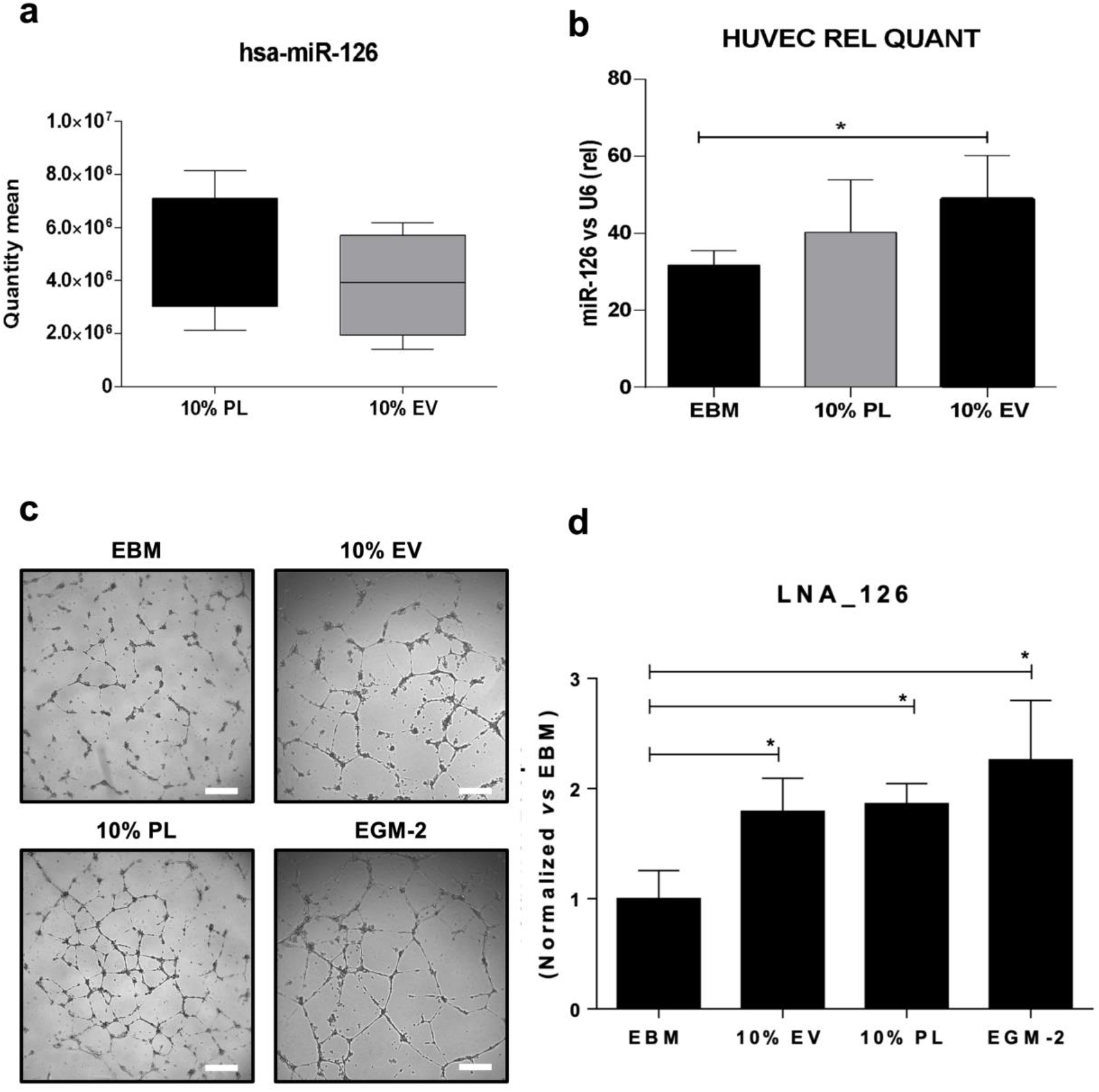
Evaluation of hsa-miR-126 in endothelial cells after treatment with PL-derived EVs. (A) Quantitative Real Time PCR, showing a similar amount of has-miR-126 among the percentages of EVs (5, 10 and 20%) and 10% PL when employed as stimulus. One way ANOVA test was applied. A range of N=3-16 experiments was performed. (B) Relative expression of has-miR-126 in HUVEC by real Time PCR after treatment with 5, 10% and 20% EVs or 10% PL. The fold change on the basal EBM is indicated. *p<0.05. (C) Representative optical images and (D) quantification of the matrigel assay of HUVEC after antagomir-126 (LNA_126) transfection and different treatments. Magnification 4X. White scale bar, 200μm. *p<0.05. (E) Targetscan analysis at the human 3’ UTR of VEGF, highlighting that it is a target of miR-126-3p in the poorly conserved category and the validation ^b^y RTPCR showing no modulatory effect of this gene in our system. A range of N=3-7 experiments was performed.

Finally, we tested the functional effects of blocking EV-miR-126 on angiogenesis and if HIF-1αor VEGF, known to upregulate miR-126^76^ and to be abundant in PL^2,46–49^, respectively, might be downstream targets. In physiological conditions matrigel assay showed that the number of loops significantly increased in presence of LNA-126 but with treatment with 10% EV or 10% PL compared to control EBM (Fig. 6c-d, all p<0.05 and supplementary Fig. 1c-d for Matrigel with Control), therefore confirming the angiogenic effect contained in the EVs derived from PL and mediated at least partially by miR-126. The analysis of the matches to human 3’ UTRs of both HIF-1α and VEGF trough the Targetscan software, revealed that only VEGF is a target of miR-126-3p in the poorly conserved category (Fig 6e). Nevertheless, the validation by real time PCR showed no modulatory effect in our system ascribable to VEGF (Fig. 6e) suggesting different molecular targets controlled by miR-126.

## DISCUSSION

This study demonstrates that PL formulations are enriched of EVs, which are biologically active and efficiently sustain angiogenesis in endothelial cells. Interestingly, the EV concentration in our PL is very high and mainly composed by a small EVs subset (50-200 nm)^58^, suggesting that PL might be reinterpreted as an abundant biological source of this EVs subpopulation.

Intriguingly, in line with reports showing comparable *in vitro* and *in vivo* haemostatic properties of both platelet microparticles and PL-derived EVs on platelets^77–79^, our EVs only partially retain this feature. In fact, our PL-derived EVs do not promote platelet aggregation, but they are able to preserve coagulation in a physiological manner, strengthening their versatile use as angiogenic/antiaggregant mean for cardiovascular applications (where platelet aggregation is a critical risk factor^80^), as a topical product for bleeding during surgery, or as a substitute of platelet transfusion. Antiaggregant therapies are acknowledged to negatively impact coagulation, causing bleeding^81^, therefore, the possibility to investigate and distinguish these two interconnected haemostatic properties in platelet-derived products (such as PL) is of paramount significance, in order to develop novel products with a unique haemostatic function compatible with the clinical use.

Our results demonstrated that EVs after internalized by endothelial cells, positively enhances angiogenesis by fostering endothelial tubule-like networks in a complex 3D microenvironment, similarly to PL. This phenomenon is in line with those describing the angiogenic effects of EVs derived from circulating intact platelets^82,83^, or from other non-platelet cell types^84,85^. Although PL also contains a plethora of several soluble mediators with angiogenic function^2,46–49^, it is conceivable that EVs implement this property that PL normally possesses. Notably, 10% EVs revealed as the optimal condition in culture, showing a non-canonical dose-response of EVs without additional effects at higher percentage in line with the variable biological effects of EVs already verified^86^. The 10% EVs might represent a sort of “balanced” amount. We have already experienced that the 10% PL itself is the optimal percentage for angiogenic assay also in presence of inhibition of specific soluble factors within the preparation. Below or above this threshold angiogenic effects are not optimal^46^. Nevertheless, some biological differences exist between PL and EVs: these latter preserve the physiological levels of hydrogen peroxide (<100 μM)^87^ more efficiently than PL and other percentages of EVs in parallel to a comparable redox status (Nox4), therefore generating an antioxidant microenvironment, known to foster beneficial angiogenic effects in endothelial cells^68,87–89^ and to preserve vascular function, homeostasis, and integrity of the vascular network beyond pathological scenarios^90^. This phenomenon also occurs in platelets: H_2_O_2_ enhances their activation^91^ or aggregation upon specific agonists^92^, resulting in a loop of specific NADPH which act as a sensor of H_2_O_2_ axis in endothelial cells^68,93,94^. This result is consistent with the lower production of H2O2 found between EVs and PL. It is plausible that EVs contain the machinery for both oxidant and antioxidants molecules, therefore acting in a double fashion according to metabolic needs and signals within the microenvironment^95^.

To date, the molecular mechanism by which clinical preparations obtained from platelets enhance regenerative angiogenesis, remains not fully explored. Both platelets, the main contributor of miRNAs released in the blood, and their microparticle counterparts, contain a wide range of overlapping miRNAs^96^, whose investigation so far has been restricted mainly to physiological functions related to aggregation and activation^96,97^. The miRNAs derived from EVs of platelets origin are both novel biomarkers in the context of anti-platelet therapies and platelets function^98^, and biological mediators in the cellular microenvironment^6,99,100^, suggesting their extra-platelet role beyond hemostatic properties.

We have demonstrated that half of the small RNA content of PL is composed by miRNAs. We found that angiogenic miRNAs (miR-320, 25 and 126) are contained in the EV cargo as well as in PL. So far, a proper screening of the miRNA profile has been performed only in PRP^101^ and intact or hyperreactive platelets from healthy subjects, or in presence cardiovascular pathologies. Interestingly, when we profiled our PL, data have shown that the formulation reflects a similar repertoire of mature miRNAs found in human platelets and described in the literature^71,72^, including the abundant miRlet-7 (a marker of platelet differentiation and maturation in megakaryocytopoiesis^102^), or defined microRNA families (*i.e*.miR-25 and 103)^72^. By intertwining the transcriptomic expression profile with the vascular function, we have confirmed that in PL-based preparations the miRNAs quantitatively more represented are also strictly interconnected to the angiogenic function. The EVs contained in PL mirror this picture and confirm that miR-126 is the most significant miRNA in both preparations.

The angiomiRNA miR-126 is one of the most abundant miRNA expressed in platelets^103,104^. Sharing with miR-320 (that we also found as highly represented) the unique expression also in endothelial cells, miR-126 is able to downregulate adhesion molecules (e.g. VCAM-1) upon influence of specific cytokines (*i.e*. VEGF), therefore contributing to endothelial migration, proliferation, activation, and vascular inflammation^75^. Exosomes enriched in miR-126 are strictly correlated with protection from ischemic events^105^, and atherosclerosis progression^106^. Changes in circulating levels of miR-126 have been described in patients with acute ischemic stroke^107^, coronary artery disease, or type 2 diabetes^108^. Moreover, vascular development and integrity are sustained by miR-126 in zebrafish and mice^109,110^, whereas *in vivo* silencing of miR-126 impaired angiogenesis^111^ upon ischemic insult. Thus, miR-126 appears as a potential biomarker and therapeutic target for angiogenesis. Nonetheless, the transfer of miRNAs in the form of EVs under physiological and pathological conditions from platelets to endothelial cells (and vice versa), and the modality by which biological functions are sustained, is still under intensive investigation.

Our data confirm that miR-126 of platelet origin plays a key role in the angiogenic homeostasis of endothelial cells. Accordingly, results highlight that HUVECs increase intra-cellular levels of miR-126 upon stimulation with EVs of PL origin, by adding exogenous miR126 by EV transfer when the endogenous miR-126 is silenced. Thus, our results demonstrate that a fraction of the angiogenic effect induced by the whole PL preparation is directly ascribable to the EV cargo, and specifically to platelet-derived miR-126.

This study has some limitations. Although we found that VEGF is a target of miR-126, we couldn’t observe any modulatory effect in our system, suggesting that alternative mechanisms are needed to be verified. Only few of them have been already described. For instance, the DNA methyltransferase (DNMT), playing a role in hypoxia tolerance, has been found as a target of miR-126 contained in exosomes^112^. Further mechanisms can coexist, including the reduction of cell apoptosis ^105^, the overactivation of autophagy trough Beclin-^113^, the novel delivery system by apoptotic bodies trough CXCL12^114^, or the inhibition of the negative regulators of the VEGF-axis^110,115^. More importantly, the angiogenic properties of both EVs and PL cannot be explained uniquely by the miR-126. The miRNome here described suggests the presence of additional miRNAs with similar function. For instance, the exosomal derived-miR25 has been found to promote angiogenesis, vascular permeabilization, metastatic niches in cancer and involved in cardiovascular disorders^116,117^.

To date, the individual contribution of EVs within PL has not been fully elucidated in terms of regenerative angiogenesis. Certainly, the methodology to manufacture PL severely impacts the quantity and the quality of EVs within the formulations and in particular that employed to concentrate, lyse or activate platelets^118^ in the formulation. Accordingly, PL preparations with excessive heterogeneity of EV content might result in parallel different downstream signaling and pathways activated with a wide range of unpredictable biological effects, also depending on cells potentially targeted by clinical PL preparations.

In conclusion, PL-based formulations are a source of both biologically available miRNAs and EVs defining the hallmark of platelet origin. The EVs reflect the “angiogenic physiology” of PL, confirming that a cell-free therapy approach may be a novel effective strategic tool in clinical applications.

Future investigations will be required to unveil the role of downstream targets of different miRNAs potentially preserved in EVs of platelet origin, and how additional processes not limited to angiogenesis are modulated, including immunomodulatory functions and paracrine effects.

## ACKNOWLEDGMENTS

We thank the Sapienza Department of Medical Surgical-Sciences and Biotechnologies in Latina for the continuous support and effort. We also thank the FlowCore in Mannheim (Medical Faculty Mannheim, University of Heidelberg) whose Cytek^®^ was funded by the Federal Ministry of Education and Research (BMBF) and the Ministry of Science Baden-Württemberg within the framework of the Excellence Strategy of the Federal and State Governments of Germany.

## AUTHORS CONTRIBUTION

A.B. performed main experiments, M. C., M.M. and R. R. 3D constructs, F. P. QPCR of small RNA seq, M. I. and E. S. isolated the EVs, O. F. and G. M. performed NTA and cytofluorimetry, X. D. developed the matrix for the angiogenic network of the small RNA seq, E. D. M. performed the droplet digital PCR, A. D. and F. P. all experiments on aggregation and oxidative states, S. M. performed the TEM, V. P. the cell transfection, G. F. and I.C. reviewed and edited the manuscript; E.D.F. conceived the study and wrote the paper.

## SUPPORTING INFORMATION

Supplementary Figure 1A-D are available in the online version of the manuscript

## DATA AVAILABILITY

Main data generated or analysed during this study are included in this article, and detailed data are available from the corresponding authors on reasonable request.

## FIGURE LEGENDS

**SUPPLEMENTARY FIGURE 1.**
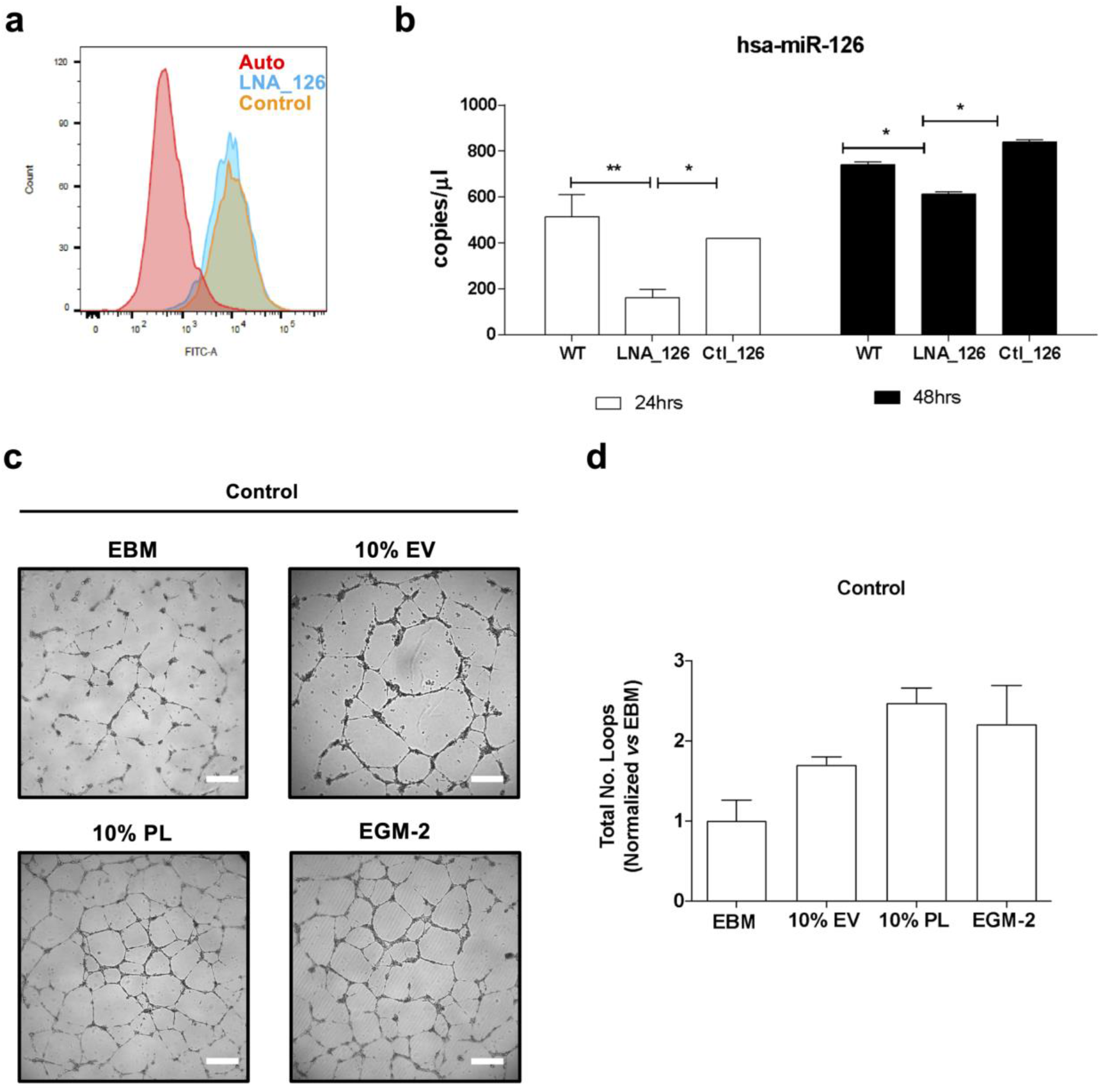
Validation of cell transfection with antagomir-126 (LNA_126) in endothelial cells. (A) Representative histogram of the cytometric analysis of HUVEC with LNA_126 (light blue line) and Control (orange line) in FITC channel. Autofluorescence is highlighted with the red line. (B) Absolute quantification of hsa-miR-126 by Droplet digital PCR, showing the efficient downregulation of the number of the copies in HUVEC after transfection with LNA_126 LNA or Control (Ctl_126) compared to untreated cells (WT) at 24h and 48h. (C) Representative optical images and (D) quantification of the matrigel assay of HUVEC after Control (Ctl_126) transfection and different treatments. Magnification 4X. White scale bar, 200μm. *p<0.05.

